# Loss of graded enemy recognition in a Whitehead population allopatric with brood parasitic Long-tailed Cuckoos

**DOI:** 10.1101/2020.02.21.958017

**Authors:** Shelby L. Lawson, Nora Leuschner, Brian J. Gill, Janice K. Enos, Mark E. Hauber

**Affiliations:** Department of Animal Biology, University of Illinois at Urbana-Champaign, Urbana, Illinois 61801, USA; School of Biological Sciences, University of Auckland, PB 92019, Auckland, New Zealand; Auckland War Memorial Museum, PB 92018, Auckland, New Zealand

**Keywords:** brood parasitism, host-parasite interactions, model presentations

## Abstract

Many avian hosts of brood parasitic birds discriminate between different types of threats and may respond with categorically different, specifically anti-predatory or anti-parasitic behaviors. Alternatively, hosts may adjust their responses to threat level in a graded manner, responding more aggressively to brood parasites during the laying and incubation stages of nesting, when nests are most susceptible to parasitism, and more aggressively to nest predators during the nestling and fledgling stages when predation would be more costly than parasitism. In New Zealand, endemic host Whiteheads act inconspicuously around their nests in the presence of sympatric Long-tailed Cuckoos, their obligate brood parasite, perhaps to avoid disclosing nest location. We tested behavioral responses of a Whitehead population on Tiritiri Matangi Island that has been breeding allopatrically from cuckoos for 17 years. We presented models of the allopatric parasite, a sympatric predator (Morepork Owl), and a sympatric non-threatening heterospecific (Song Thrush) during the egg and chick stages, and to groups of cooperatively breeding Whiteheads. We compared responses across nest stage and stimulus type. We found that, unlike sympatric Whiteheads elsewhere in New Zealand, Whiteheads on Tiritiri Matangi produced alarm calls in response to the cuckoo model. Furthermore, the rate of alarm calling was similar towards the cuckoo and the owl and across egg and chick stage and higher than to the control stimulus. These results are consistent with allopatric Whiteheads having lost their specific anti-parasitic defense tactics in response to brood parasitic cuckoos.

## INTRODUCTION

Obligate brood parasitism is costly for host species that are left to care for the unrelated young at a cost to themselves and their own young (e.g. Payne 1977, Røskaft et al. 1990). To reduce the costs of brood parasitism, effective antiparasitic strategies include host species removing parasitic eggs or nestlings, or abandoning parasitized clutches altogether (Davies 2000, Soler, 2017). However, some host species cannot discriminate parasitic eggs or chicks from their own, and thus face strong selection to prevent parasitism in the first place (McLean and Maloney 1998, Feeney et al. 2012). Accordingly, hosts may respond with high levels of aggression towards adult brood parasites near their nests in order to deter parasitism (e.g. Davies and Welbergen 2009, Yasukawa et al. 2016). Nest defense, however, in itself can be a costly behavior that reduces time that could be used for foraging or feeding young (Ueta 1999). Thus, hosts should only respond when actual threat is posed to the nest (Neudorf and Sealy 1992), and to an extent positively related to the current value of nest content (Regelmann and Curio 1983, Campobello and Sealy 2010).

Hosts of brood parasites may be able to discriminate between different kinds of threats when responding specifically in an anti-predatory or anti-parasitic manner to different threats at their nests, and may not respond at all to innocuous stimuli (Grim 2005). Several studies using stuffed, wooden, or other 3D model stimulus presentations have found that hosts are indeed able to discriminate brood parasites from adult predators and nest predators, and those from innocuous controls (e.g., Burgham and Picman 1989, Duckworth 1991, Neudorf and Sealy 1992, Grieef 1995, Gill and Sealy 1996, Welbergen and Davies 2008, Campobello and Sealy 2010, Trnka and Prokop 2012, Henger and Hauber 2014). Hosts of brood parasites that can discriminate between brood parasites and nest predators may be able adjust their responses depending on the threat level posed by the stimulus type; for example, models of brood parasites, presented during the laying or incubation stages of nesting (when nests are more susceptible to costly parasitism), elicit more aggression from some hosts relative to when the same model is shown during nestling or fledgling stage (e.g. Mclean 1987, Hobson and Sealy 1989, Neudorf et al. 1992). In contrast, many hosts respond to models of nest predators throughout all nest stages, sometimes more intensely with increasing age of the nest, as the contents already represent greater reproductive value with the more advanced nesting stages (e.g. Regelmann and Curio 1983, Moksnes et al. 1991, Neudorf et al. 1992, Campobello and Sealy 2010, Li et al. 2015).

Some host species are known to respond to brood parasites with unique, specific, and nonaggressive behaviors that also change throughout the nesting cycle. During laying and incubation, for example, Yellow Warblers *Setophaga petechia* utter functionally referent “seet” calls in the presence of Brown-headed Cowbirds *Molothrus ater*, a common brood parasite, that prompts nest defense in female warblers (Hobson and Sealy 1989; Gill 1995). This response is rarely seen in parents with nestlings, indicating its specificity to parasitic cowbirds and the higher risk of brood parasitism during early nesting (Hobson and Sealy 1989, Gill 1995). New Zealand’s endemic Whiteheads *Mohoua albicilla* are another species that respond in a unique way to models of their obligate brood parasite, the Pacific Long-tailed Cuckoo *Urodynamis taitensis* (Keast 1976; Gill and McLean, 1986). Specifically, McLean (1987) found that when Whiteheads, on the island of Hauturu (Little Barrier Island), New Zealand, were presented with the sympatric cuckoo’s model during early stages of nesting, they quietly returned to their nests to guard the content. This secretive behavior is thought to reduce the possibility of the cuckoos’ discovery of the host nest, because loud conspicuous behaviors can inadvertently reveal the location of a nest (e.g., alarm calling) (Banks and Martin 2001). During the nestling stage, however, Whiteheads responded aggressively to the cuckoo model with alarm calling, and more individuals responded in general (McLean 1987). Thus, Whiteheads adjusted their behavior in accordance with the threat posed by the cuckoo, and the increased value of their nest contents over time.

Responses to brood parasites vary not only by nest stage but also by geographic overlap with the parasites, such that host species breeding apart from brood parasites (allopatry) have often been found to lack parasite-specific responses and/or parasite-predator discrimination (Røskaft et al. 2002). For example, Yellow Warblers breeding allopatric from Brown-headed Cowbirds in Alaska, when presented with a cowbird model, rarely produce anti-parasitic seet calls, and they display similar aggression towards the cowbird model as to innocuous controls, indicating that recognition of cowbirds as a threat has been lost (Briskie et al. 1992; Kuehn et al. 2016).

In addition to the North Island and Hauturu of New Zealand, Whiteheads also occur on Tiritiri Matangi Island in the Hauraki Gulf, where they were translocated from Hauturu in 1989 (Rimmer, 2004) and have since been breeding allopatrically from Pacific Long-tailed Cuckoos (Marchant et al. 1990). Thus, Whiteheads present an opportune system with which to test whether hosts that have specific anti-parasitic behaviors in sympatry retain these behaviors (and therefore enemy recognition of brood parasitic cuckoos) after a relatively short period of allopatry relative to the maximum lifespan (16 years) of individuals in this species (Leuschner et al. 2007). Using a model exposure experiment, we compared aggressive responses (alarm calling and mobbing) towards cuckoos to those towards the adult- and nest-predatory Morepork Owl (*Ninox novaeseelandiae*) at known nest sites and group nesting sites. The Morepork is a nocturnal predator native to Tiritiri that occasionally prey on small passerine adults and nestlings (Haw et al. 2001), and these owls can be seen being mobbed by small passerines during the day (Falla et al. 1979). We predicted that if Whiteheads retained their anti-parasitic defenses towards cuckoos (quiet movement away in the presence of cuckoo), then we would see stronger aggressive responses towards the Morepork than the Pacific Long-tailed Cuckoo. We also examined responses at known nest sites across the Whiteheads’ nesting stages, and predicted that aggressive responses would be stronger to the cuckoo during the nestling stage compared to egg stage similar to McLean’s (1987) findings, because whereas anti-parasitic sneaking behavior is most crucial during egg stage when nests are at the highest risk of brood parasitism, aggressive defenses are more pertinent in nestling stage when cuckoos represent nest predatory risk (Beaven, 1997, Gill et al. 2018). We used a model of a Song Thrush (*Turdus philomelos*) as an innocuous sympatric control species (Figure 1).

**Fig. 1.**
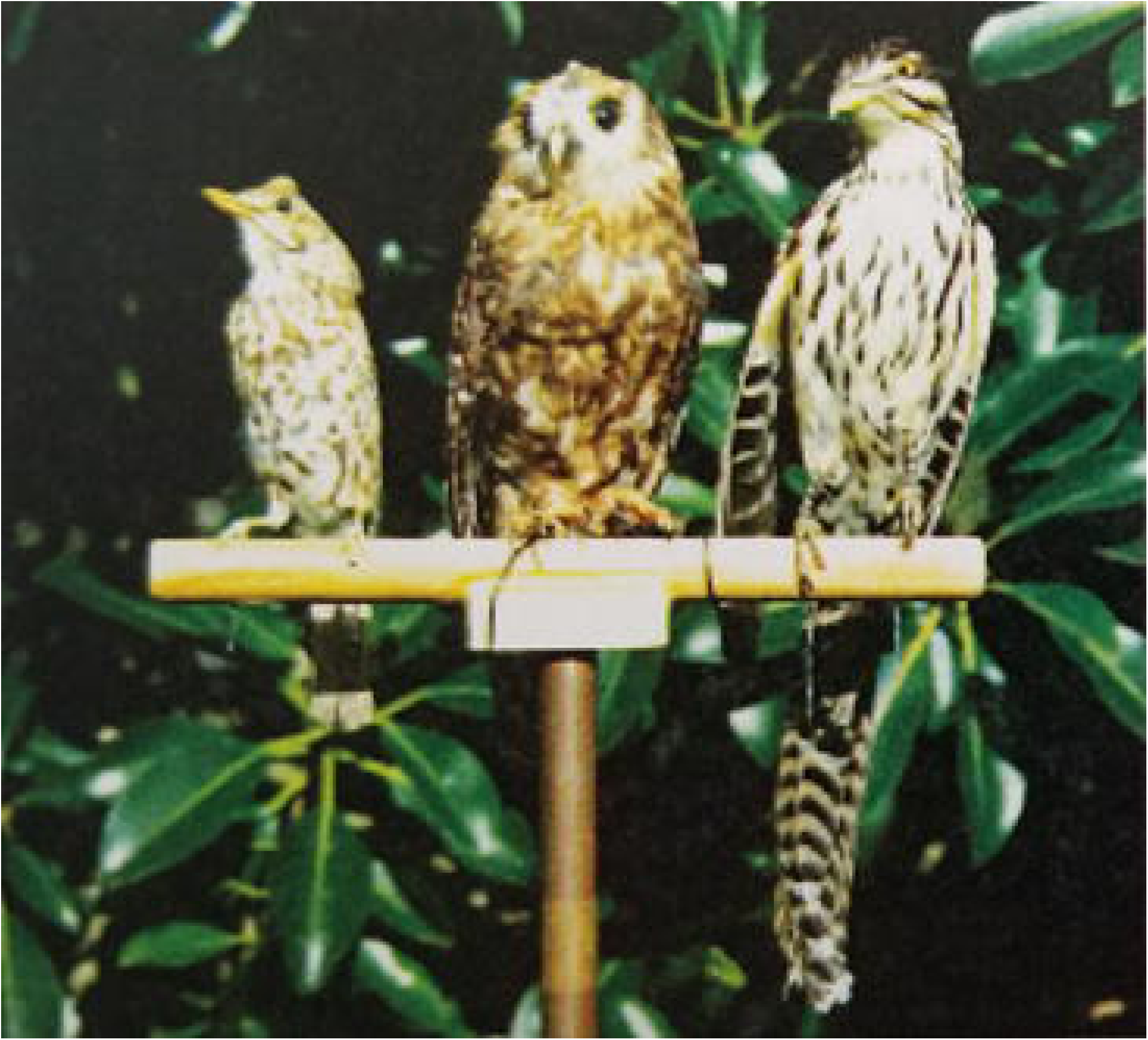
Image of models used for the study. From left to right: Song Thrush, Morepork Owl, Long-tailed Cuckoo.

## METHODS

We studied Whiteheads on Tiritiri Matangi Island, Hauraki Gulf, New Zealand. For 100 years Tiritiri was used as farmland, thus 90% of its native bush had been cleared (Rimmer, 2004). In 1980, Tiritiri became a Scientific Reserve, and a plan was established to re-establish native flora and fauna (Drey et al. 1982). At that time the island was mostly covered with scrub, grassland and fern, but within 10 years 280,000 trees were planted by volunteers (Rimmer 2004). Most land mammals, all of which are introduced species in New Zealand, were eradicated during this period, although the Pacific rat, kiore *Rattus exulans*, was not removed until 1993 (Rimmer 2004). Bird translocations, including two releases of Whiteheads captured on Hauturu, took place in 1989 (40 Whiteheads) and 1990 (40 Whiteheads) (Rimmer 2004). Though there is a large, annually breeding population of Whiteheads on the island, Pacific Long-tailed Cuckoos do not breed on the island, thus Whiteheads on Tiritiri have been breeding in allopatry from their brood parasites for 17 years prior to this study (Marchant et al. 1990). Whiteheads are cooperative breeders that often form groups with one or two breeding pairs and several secondary helpers (often related to the breeding pair/s), but will occasionally breed in single pairs as well (McLean and Gill 1988, Gill and McLean 1992); Whiteheads are one of the only species parasitized by the Long-tailed Cuckoo on and off the North Island of New Zealand, and both groups and single pairs are susceptible to parasitism.

For this study, we began to visually search the vegetation on island for active Whitehead nests during the austral spring in September 2006, and continued throughout the experiment. Models were presented at known nests and at cooperative group sites presumed to have nests from October-December, 2006. Using a stuffed avian model-presentation approach (recommended in Sealy et al. 1998, Grim 2005), we presented three different taxidermic models consisting of: a Pacific Long-tailed Cuckoo (hereafter: cuckoo) model was used as brood parasite threat, a Morepork (hereafter: owl) model was used a predatory threat (for both nests and adult Whiteheads), and a Song Thrush was used as an innocuous sympatric control species. We used only one model per species as this was all that was available to us from the Auckland Museum. Presumably Whiteheads have experience with the owl and the Song Thrush as both inhabit Tiritiri Matangi Island (Heather and Robertson 2000).

Trials were run near known nests as well as sites where groups of Whiteheads were detected but nests could not be located. Trials were conducted either during laying and early incubation (egg stage), or when chicks were about 10 days old (chick stage). Nests were difficult to find and limited in numbers. Thus, the same nests underwent separate trials for all three models during both the egg and chick stages (if available), presented in a randomized order to prevent order effects. The group sites were also tested with each model in random order on different days. Group sites were far enough apart to safely assume no group was tested twice with the same model type. During trials we quantified alarm calling and mobbing responses to models, both of which are considered general aggressive behaviors.

Before starting a trial, the preferred flight path of nest owners was observed so that the model could be placed in the most visible location. Groups were observed before trials to determine an active location to place the model. After the breeding group had left the vicinity of the nest, one of the models was placed at a lateral distance of 2 m from the nest. Models were fastened to metal pipes attached to a tripod that was 2.10 m off the ground. All models were placed facing the nest. Trials ran for 5 minutes each after one or more Whiteheads responded. If Whiteheads did not respond within 15 minutes, the trial was terminated.

We quantified alarm calling by categorizing alarm call rate per 30 second intervals over 5-minutes and then taking the mean of the score: (no vocalization = 0, 1-10 calls/interval = 1, 11-20 calls/interval = 2, 21-30 calls/interval = 3). We used this method due to the high rate of alarm calling, making it difficult to count and calculate and actual call rate (see Grim 2005). We quantified mobbing by counting the number of Whiteheads that alarm called to a model (abundance).

Six nests were found and tested along with 13 group sites. Two nests failed to produce nestlings and thus were only tested at the egg stage. Two nest site trials could not be conducted due to inclement weather. Thus, in total we conducted 24 trials at nest sites with the cuckoo (n = 5 egg; n = 4 nestling), owl (n = 4 egg; n = 4 nestling) and thrush models (n = 4 egg; n = 3 nestling), and 38 total trials at group sites with the cuckoo (n = 13), owl (n = 13), and thrush models (n = 12).

### Statistical Analyses

We first evaluated how model type affected our response variables of interest (call rate and abundance) with data from all sites, using a separate model for each response variable in SAS/STAT software 9.4, SAS Institute Inc., Cary, NC, USA. We used general linear mixed models to analyze call rate, and a generalized linear mixed model fitted with a Poisson distribution for abundance as data were not normally distributed. We included model type as a main effect and group ID as random factor. We initially included trial order as a main effect, but removed the term as it was not significant in either model (*F*_5,36_ = 0.53-1.03, *P* = 0.42-0.76).

We then examined if responses to model type varied by nest stage (egg versus chick) using data from only nest sites because nest stage was known. We used the same statistical model structures per response variable described above, but included model type, nest stage, and a model type*nest stage interaction as main effects. We again included trial order as a main effect initially, but it was not significant, and was removed (*F*_5,9_ = 1.51-2.45, *P* = 0.11-0.27).

Permits for this study were obtained from the New Zealand Department of Conservation and approved by the University of Auckland Ethics Committee. No live birds were handled during this experiment.

## RESULTS

### Responses of Whitehead at nests and in groups

Call rate significantly differed in response to model types (*F*_2,41_ = 22.49, *P* < 0.001) (Fig. 2). Based on post-hoc pair-wise comparisons of least-squares means, Whiteheads called less frequently at the Song Thrush compared to the cuckoo (*F*_2,41_ = 9.93, *P* < 0.001) and owl model (*F*_2,41_ = 9.49, *P* < 0.001), but call rate did not differ between the cuckoo and owl treatment (*F*_2,41_= 0.16, *P* = 0.43). There was a significant difference in the number of Whiteheads responding to model types (*F*_2,41_ = 17.82, *P* < 0.001) (Fig. 3). Based on post-hoc pair-wise comparisons of least-squares means, more Whiteheads responded during trials with owl *(F*_*2*,41_ = 9.63, *P* < 0.001) and cuckoo (*F*_2,41_ = 9.02, *P* < 0.001) models compared to Song Thrush, but there was no statistical difference between the number of Whiteheads responding to owl and cuckoo trials (*F*_2,4 1_= −0.51, *P* = 0.69).

**Fig. 2.**
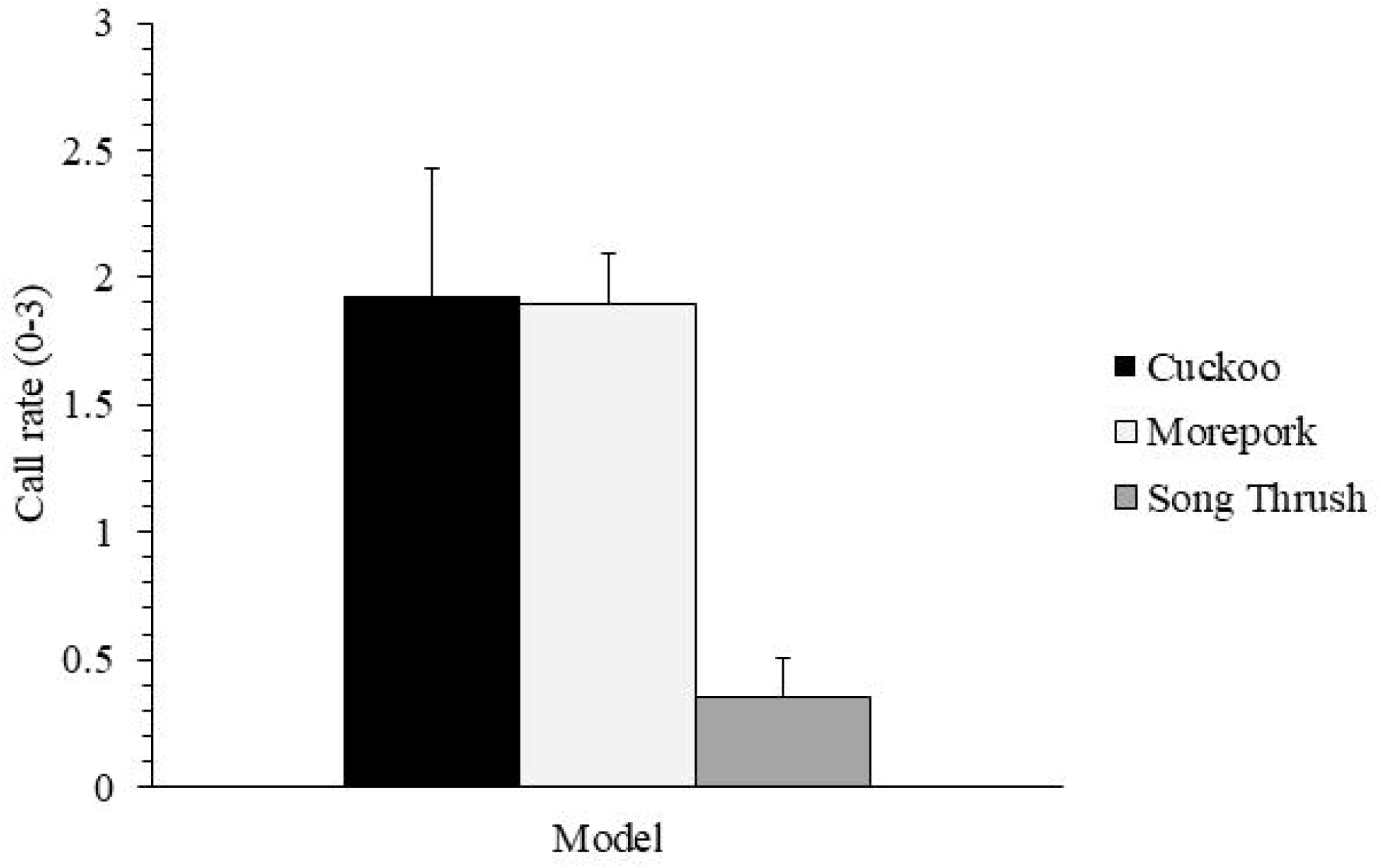
Call rate (mean (±SE; measured in discrete categories 0-3) of all responding Whiteheads during 5 minute trials at group and nest sites combined. Call rate was scored by the frequency of alarm calls per 30 second intervals (no vocalization = 0, 1-10/interval = 1, 11-20/interval = 2, 21-30/interval = 3).

**Fig. 3.**
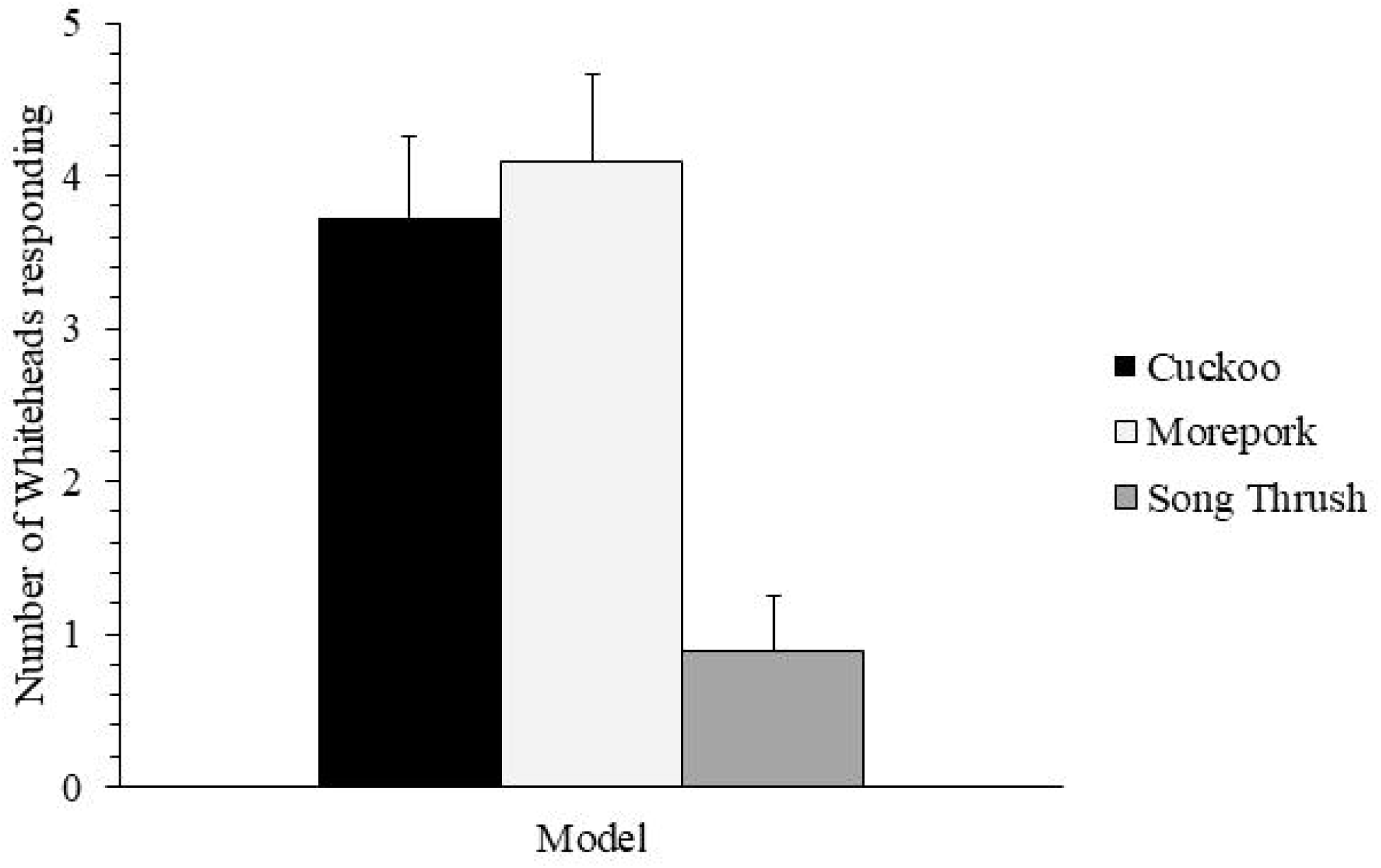
Number of Whiteheads (mean±SE) responding during 5 minute trials at group and nest sites combined.

### Effect of nest stage on responses

As shown in the previous statistical outputs which pooled the group and nest sites together, we found that at known nest sites, the call rate differed significantly between model types (model treatment term: *F*_2,13_= 3.53, *P* = 0.05) (Fig. 4). Nest stage, however, did not significantly affect call rate (nest stage term: *F*_1,13_ = 0.91, *P* = 0.35) and there was no statistically significant interaction between model and nest stage (model type*nest stage term: *F*_2,1 3_= 0.46, *P* = 0.64).

**Fig. 4.**
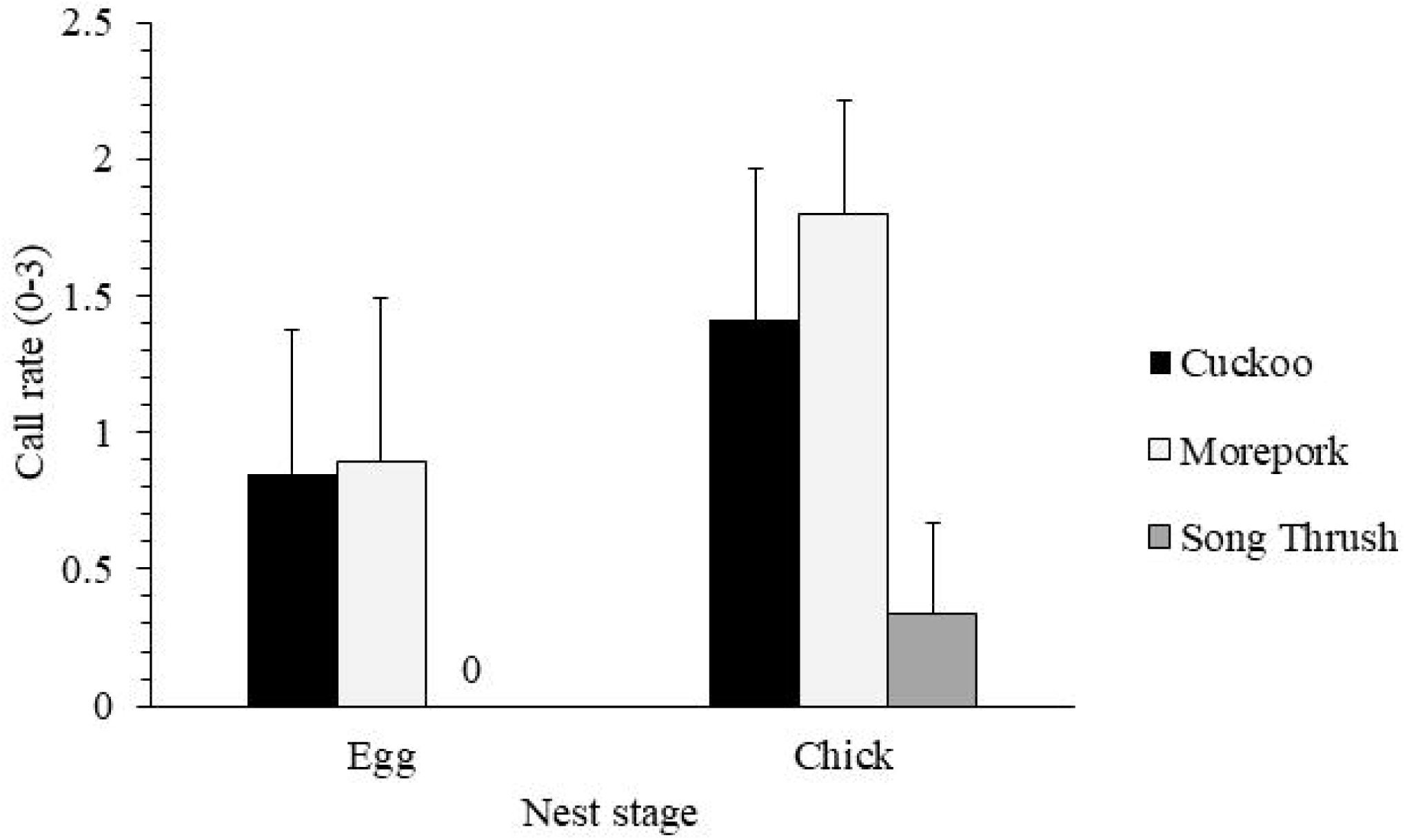
Call rate (mean±SE) of all responding Whiteheads during 5 minute trials at nest sites. Call rate was scored by the frequency of alarm calls per 30 second intervals (no vocalization = 0, 1-10/interval = 1, 11-20/interval = 2, 21-30/interval = 3)

In contrast to the findings from the sites combined, the number of Whiteheads responding did not significantly differ between model treatments at nest sites (*F*_2,13_ = 2.72, *P* = 0.103) (Fig. 5). Nest stage, however, affected how many Whiteheads responded to model types (*F*_1,13_ = 7.67, *P* =0.015) as more individuals responded during chick stage than egg stage (Fig. 5). Finally, there was no statistically significant interaction between model type and nest stage (*F*_2,1 3_= 2.04, *P* = 0.170).

**Fig. 5.**
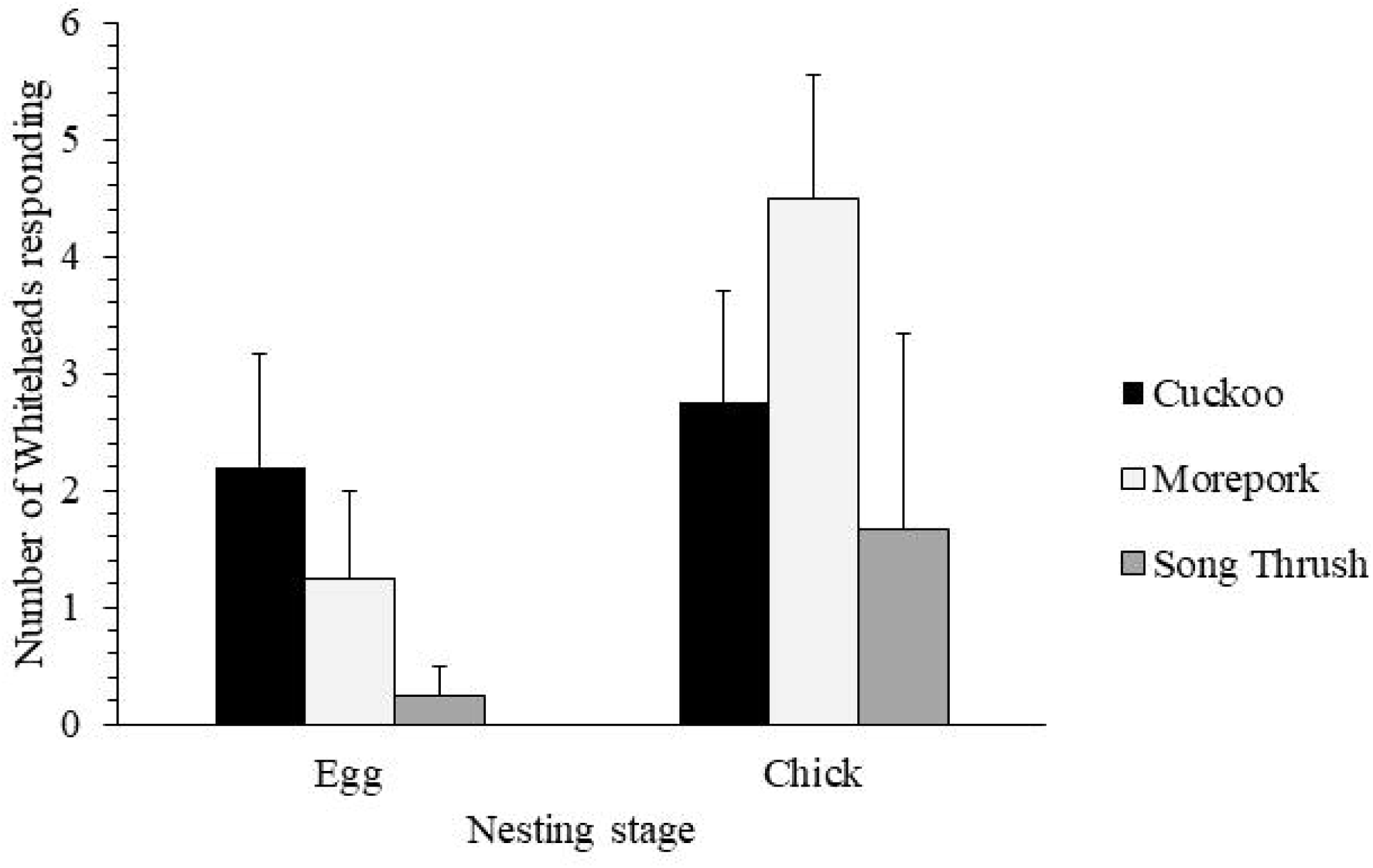
Number of Whiteheads (mean±SE) responding during 5 minute trials at nest sites.

## DISCUSSION

Our data showed that Whiteheads on Tiritiri Matangi Island can discriminate and respond accordingly to the perceived threat class of the species when comparing models of predators vs. innocuous species. At nest and group sites combined, the models that were discovered by Whiteheads generated quantitative differences in the birds’ reaction to the different stimulus types. Whiteheads were particularly aggressive towards both the owl and cuckoo models, and started alarm calling as soon as the respective model was spotted. On the other hand, Whiteheads paused and noted the presence of the Song Thrush, but then resumed their usual behavior. Whiteheads viewed both the cuckoo and owl as threats, and responded with similarly high levels of alarm calling to both of these species. However, alarm calling rates towards the models were not affected by nesting stage.

### Lost defense mechanisms in allopatry

Critically, the behavior of Whiteheads towards Long-tailed Cuckoo models in allopatry studied here is strikingly different compared to Whiteheads in sympatry tested by McLean (1987). Using a similar setup to ours with model presentation trials, McLean found that during early incubation, when nests are at the highest risk of parasitism, Whiteheads behaved inconspicuously in the presence of a cuckoo model, quietly returning to their nest presumably to avoid giving away its location to the cuckoo (McLean 1987). During the chick stage, by contrast, Whiteheads became aggressive and responded with alarm calls towards the cuckoo model. Long-tailed Cuckoos are known to sometimes prey on songbird eggs and nestlings (Beaven, 1997, Gill et al. 2018), thus during the chick stage cuckoos might be viewed as nest predators, because brood parasitism is no longer a severe risk to the hosts’ reproductive output. Our research shows that in allopatry from their brood parasites, Whiteheads on Tiritiri responded to the cuckoo model with alarm calls during both egg and chick stage, suggesting loss of their inconspicuous anti-parasitic behavior seen in the sympatric population.

Whiteheads on Tiritiri had been allopatric from Pacific Long-tailed Cuckoos for 17 years prior to this study (Marchant et al., 1990). Although these cuckoos are occasionally spotted on the island, there is not a breeding population; thus, our subjects tested had no experience with cuckoos as brood parasites and, presumably, no experience with them as nest predators, either. This dynamic reduces the possibility of aggression towards cuckoos as a learned behavior (Davies and Welbergen 2009), and so it appears that the Whitehead’s aggression towards the cuckoo is mediated by mechanisms that requires no previous experience.

Similar differences in response to threat types between allopatric and sympatric populations of avian hosts and brood parasites have been seen in other systems as well. Some hosts that have become allopatric from brood parasites display decreases in anti-parasitic defenses (Briskie et al. 1992; Gill and Sealy 2004; Hale and Briskie 2007), including the loss of foreign-egg rejection behaviors (Cruz and Wiley 1989; Marchetti 1992). However, egg rejection behavior has been hypothesized to be more costly to maintain than aggressive nest defense (Cruz and Wiley 1989) and, so, nest defense in allopatry would presumably be lost at a slower rate compared to egg rejection. Whiteheads on Tiritiri appear to view cuckoos as a general nest predator threat, similar to the owls, rather than a specific parasitic one, as indicated by the lack of specific anti-parasitic behaviors and the use of anti-predatory behaviors instead. The latter behaviors are likely easier to maintain than re-evolving or maintaining anti-parasitic responses (Hosoi and Rothstein 2000), and because cuckoos also represent a nest predation risk at all breeding stages, it is adaptive for Whiteheads to respond with aggression.

In the pooled nest and group data, the owl and cuckoo models attracted more Whiteheads than the Song Thrush. It is possible that the higher alarm calling rates during trials for owl and cuckoo models attracted more conspecifics to the site to mob. Accordingly, in one instance, two neighboring flocks were attracted to the area after the nest owners alarmed-called at the owl model, resulting in 12 total birds mobbing the owl. Mobbing in response to nest predators is a major fitness benefit to having neighbors or living in a group (Shields 1984, Feeney et al. 2013).

There were no significant differences in number of Whiteheads responding between nesting stage, although such lack of statistical effects may be due to our low sample sizes at known nests. The discrepancy between some of the nest-site only data and the combined nest-group site data for the model type’s impact on the numbers of Whiteheads responding may be due to the small sample of nests that could be tested compared to the larger pooled data set, but it is also possible that we saw this discrepancy because some Whiteheads at group sites were seen to have fledglings, compared to nest sites which had only eggs or chicks. Offspring become more valuable as their age increases (Regelmann and Curio 1983); thus, it is not surprising that Whiteheads at group sites respond strongly and in higher numbers to potential threats than at nest sites, causing the pooled data for number of Whiteheads to yield different statistical tests.

There are some severe limitations in our study. Due to the cryptic nature of Whiteheads and the difficulty of locating their nests, our sample sizes were small, which reduced our statistical power. We also did not capture, sex, and band Whiteheads for individual identification, making it impossible to determine any potential sex differences in response to the models. Without banding individuals, it is possible that even on Tiritiri, during the cuckoo model trials, the incubating females quietly returned to nests (as reported in sympatry), and that we only measured aggressive behaviors from males in this study. However, in every nest site cuckoo trial where there was any aggressive response, more than one Whitehead responded and produced alarm calls, which likely included the females.

Overall, after a short amount of time relative to their generation time, Whiteheads on Tiritiri do not display specific anti-parasitic responses to Pacific Long-tailed Cuckoos and exhibit a non-specific aggressive response when presented with a cuckoo model similar to responses to nest-predatory Morepork. Moreover, this response does not appear to change throughout the breeding cycle, indicating that the Whiteheads on Tiritiri do not view cuckoos as brood parasites, but likely as predatory threats like the Morepork. Our results are consistent with that anti-parasitic behaviors are costly to maintain, and may be rapidly lost in favor of cheaper general aggressive responses when there is no longer a benefit to maintaining the original type, specificity, and dynamics of defensive behaviors.

